# α-carboxysome size is controlled by the disordered scaffold protein CsoS2

**DOI:** 10.1101/2023.07.07.548173

**Authors:** Luke M. Oltrogge, Allen W. Chen, Thawatchai Chaijarasphong, Julia B. Turnšek, David F. Savage

## Abstract

Carboxysomes are protein microcompartments that function in the bacterial CO_2_ concentrating mechanism (CCM) to facilitate CO_2_ assimilation. To do so, carboxysomes assemble from thousands of constituent proteins into an icosahedral shell which encapsulates the enzymes rubisco and carbonic anhydrase to form structures typically >100 nm and >300 megadaltons. Although many of the protein interactions driving the assembly process have been determined, it remains unknown how size and composition are precisely controlled. Here we show that the size of α-carboxysomes is controlled by the disordered scaffolding protein CsoS2. CsoS2 contains two classes of related peptide repeats which bind to the shell in a distinct fashion, and our data indicate that size is controlled by the relative number of these interactions. We propose an energetic and structural model wherein the two repeat classes bind at the junction of shell hexamers but differ in their preferences for the shell contact angles, and thus the local curvature. In total, this model suggests that a set of specific and repeated interactions between CsoS2 and shell proteins collectively achieve the large size and monodispersity of α-carboxysomes.

## Introduction

Spatial organization is a fundamental feature of biology–delineating the boundaries between cells and the environment and segregating the internal metabolism.^1^ Cyanobacteria and diverse chemoautotrophic bacteria use specialized protein microcompartments called carboxysomes to provide a CO_2_ enriched micro-environment for the carboxylase rubisco, which fixes CO_2_ into sugars.^2–4^ Carboxysomes are a striking example of emergent order, with thousands of individual copies of ten different constituent proteins spontaneously assembling into quasi-icosahedral particles with well-defined size and composition.^5^ The mechanism of size control is particularly interesting, as the assembled particles extend far beyond the dimensions of any individual component. The assembly process also requires high-fidelity with potential for severe phenotypic penalty. Morphological defects to the carboxysome, such as by deletion of shell genes, break the CO_2_ concentrating mechanism and result in cell death.^6–8^

Carboxysomes occur in two evolutionarily distinct lineages: α, found in most marine cyanobacteria and several clades of bacterial chemoautotrophs; and β, found in certain freshwater cyanobacteria. β-carboxysomes tend to be larger (100-400 nm in diameter) and are thought to assemble in an inside-out manner with the formation of a dense rubisco kernel preceding encapsulation by the shell proteins.^9–11^α-carboxysomes are somewhat smaller (80-120 nm in diameter) and more regular in size. Their assembly is hypothesized to involve the concurrent formation of cargo and shells.^12–14^ In this work we focus on α-carboxysomes to understand what factors drive the size and regularity of the particles.

α-carboxysome assembly culminates in particles that are densely packed with cargo, free of shell defects to serve as a CO_2_ permeability barrier, and of uniform size with diameters of about 115 nm in the model species *Halothiobacillus neapolitanus*.^4,15^ The process by which carboxysome size is determined is not understood, though several models have been put forward.^16–18^ Structural studies on the shell proteins have revealed spontaneous assembly of small icosahedral complexes with triangulation numbers of T = 3, 4, or 9.^19–21^ By comparison, the *H. neapolitanus* carboxysome is significantly larger with T ∼ 75.^22^ This suggests that intrinsic curvature preferences of the shell proteins alone^§^ are not determinative of the carboxysome size–as is the case for many small icosahedral virus capsids–but that an interplay with internal components is ultimately responsible for particle size.

α-carboxysomes are composed of a thin icosahedral shell made up of thousands of hexameric protein capsomers (CsoS1) and pentameric capsomers (CsoS4) forming the twelve vertices.^22^ Inside the shell is a dense cargo of enzymes: rubisco and carbonic anhydrase; and the scaffolding protein CsoS2 (Fig. 1).^23–26^ CsoS2 is unique among the carboxysome proteins as it is predicted to be largely structurally disordered along its entire length.^4,13,27^ It does, however, contain several distinct repeated sequence motifs that collectively define a three-part domain structure: the N-terminal domain (NTD) with N-peptide motifs, the middle region (MR) with M-peptide motifs, and the C-terminal domain (CTD) with C-peptide motifs and a conserved C-terminal peptide (CTP).

**Figure 1.**
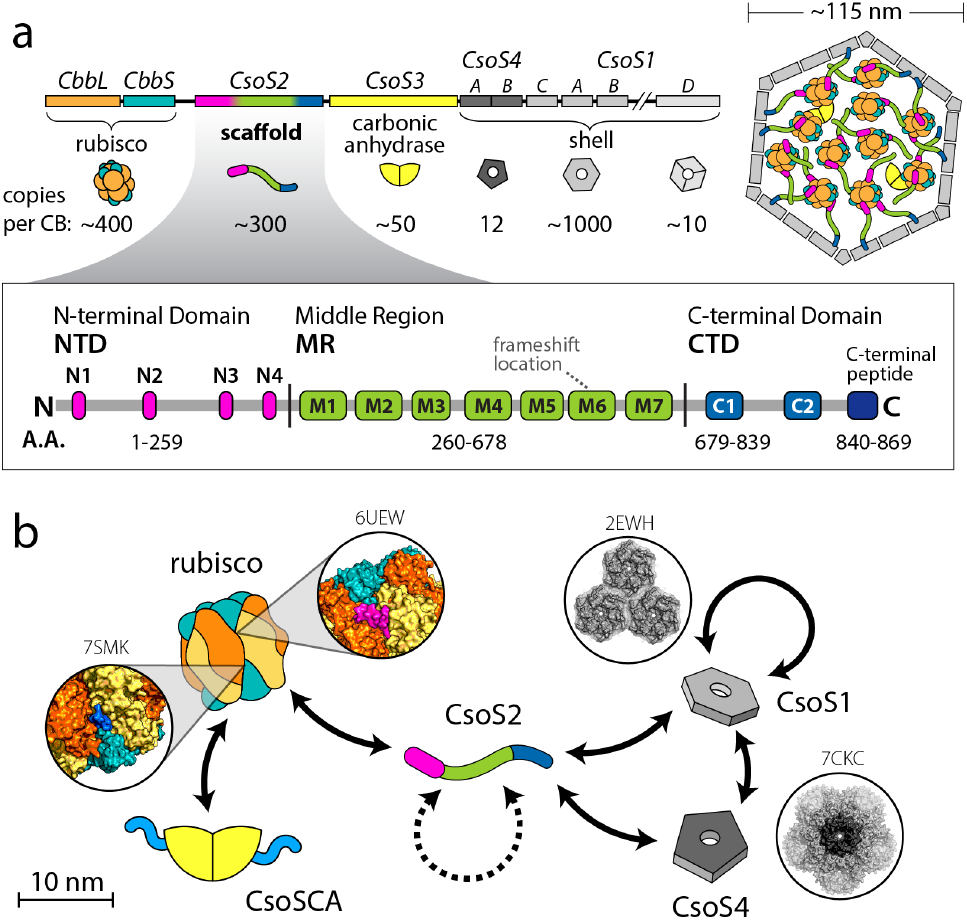
**a**) The *H. neapolitanus* regulon with the ten proteins making up the α-carboxysome. The callout shows the domain structure with repeated peptide motifs in the disordered scaffold protein CsoS2. Note that M7 has been reclassified as an M-peptide (see Fig. S1). **b**) Carboxysome protein interactome. Known protein-protein interactions are depicted with solid arrows and cases where the contact structures are solved are shown in the popouts along with their corresponding PDB IDs. CsoS2 is an important hub in the network, bridging the cargo to shell connection. Cartoon schematics are shown roughly to scale.

Previous work has shown that CsoS2 is essential to growth and acts as a central node with interactions to both rubisco and CsoS1.^13,14,28^ Specifically, the NTD binds to and encapsulates rubisco via interactions with the N-peptides,^14,25^and the CTD has been shown to associate with the shell.^20,21^ We recently showed that MR also interacts with the shell, though what distinguishes its functional role from the CTD is not yet clear.^29^

Here we describe how the sequence of the scaffold CsoS2 and, in particular, the relative numbers of M- and C-peptide motifs specify the size of the carboxysome. Furthermore, this size effect is largely independent of the primary cargo rubisco and points toward a model for size control that is driven by the shell’s mechanical properties and, specifically, CsoS2’s modulation of these mechanical properties. Finally, in light of a recent cryoEM structure of portions of the CTD-shell interface^21^ we propose a structural model of the M-peptide’s role.

## Materials and Methods

### Cloning and expression

All recombinant carboxysome expression was based on the pHnCB10 plasmid which contains the full carboxysome regulon from *Halothiobacillus neapolitanus* composed of ten proteins (Fig. 1a) under the control of a lac operator, has a ColE1 origin, and carries a chloramphenicol resistance marker.^30^ Uniprot protein IDs are as follows: CbbL (rubisco large subunit) O85040; CbbS (rubisco small subunit) P45686; CsoS2 O85041; CsoSCA (carbonic anhydrase) O85042; CsoS4A (pentameric capsomer) O85043; CsoS4B (pentameric capsomer) O85044; CsoS1C (hexameric capsomer) P45688; CsoS1A (hexameric capsomer) P45689; CsoS1B (hexameric capsomer) P45690; CsoS1D (pseudo-hexameric capsomer) D0KZ73.

For testing different CsoS2 variants and truncations, a Golden Gate destination vector was created from pHnCB10, replacing the original CsoS2 with lacZ bracketed by BsaI type II restriction sites. The different CsoS2 constructs were built by PCR amplification of the desired fragments with primers containing flanking BsaI sites and compatible recombination sites. The polyproline II helix sequence was purchased as a gBlock from IDT. All plasmids were completed by Golden Gate Assembly.^31^ The co-expression plasmids for CsoS1ABC (Fig. 2b), CsoS2A, CsoS2B (Fig. 3c), and fragments of the MR fused to GFP (Fig. S7) were built on a pFA backbone with a p15a origin, kanamycin resistance, and tetracycline operator.^32^ All constructs are summarized in Table S1.

**Figure 2.**
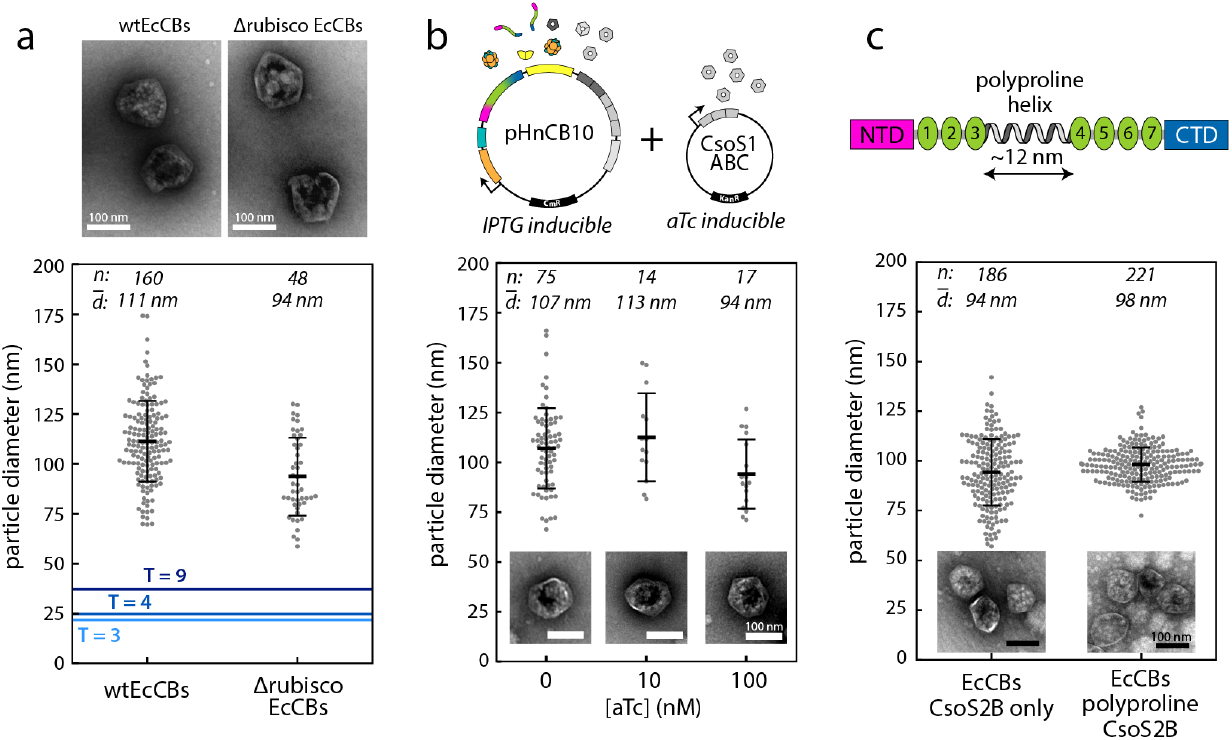
**a**) Comparison of recombinant carboxysomes with and without the primary cargo rubisco. TEM negative stain micrographs highlight the same overall pseudo-icosahedral structure. All swarm plots show the minimum Feret diameter. The dark central line is mean diameter with whiskers to +/-1 standard deviation. Particle counts and mean diameters are shown along the top. The blue lines mark the sizes of solved icosahedral complexes of shell capsomers and the corresponding T-numbers are indicated.^20,21^ **b**) Hexamer titration. *E. coli* carrying pHnCB10, a plasmid containing all ten carboxysome proteins under control of lac operator, and a plasmid with the major shell proteins CsoS1ABC under tetracycline induction control. Cells were all induced with 1 mM ITPG and variable amounts of anhydrotetracycline (aTc) to change shell expression levels. Swarm plots of particle sizes as in a) with inset zoomed TEM micrographs showing normal morphology. **c**) Polyproline helix. Carboxysomes were expressed with CsoS2B, ie. CsoS2 with no frameshift, and CsoS2B with a 12 nm polyproline II helix inserted. Swarm plots of particle sizes as in a) with inset TEM micrographs of representative particles.

**Figure 3.**
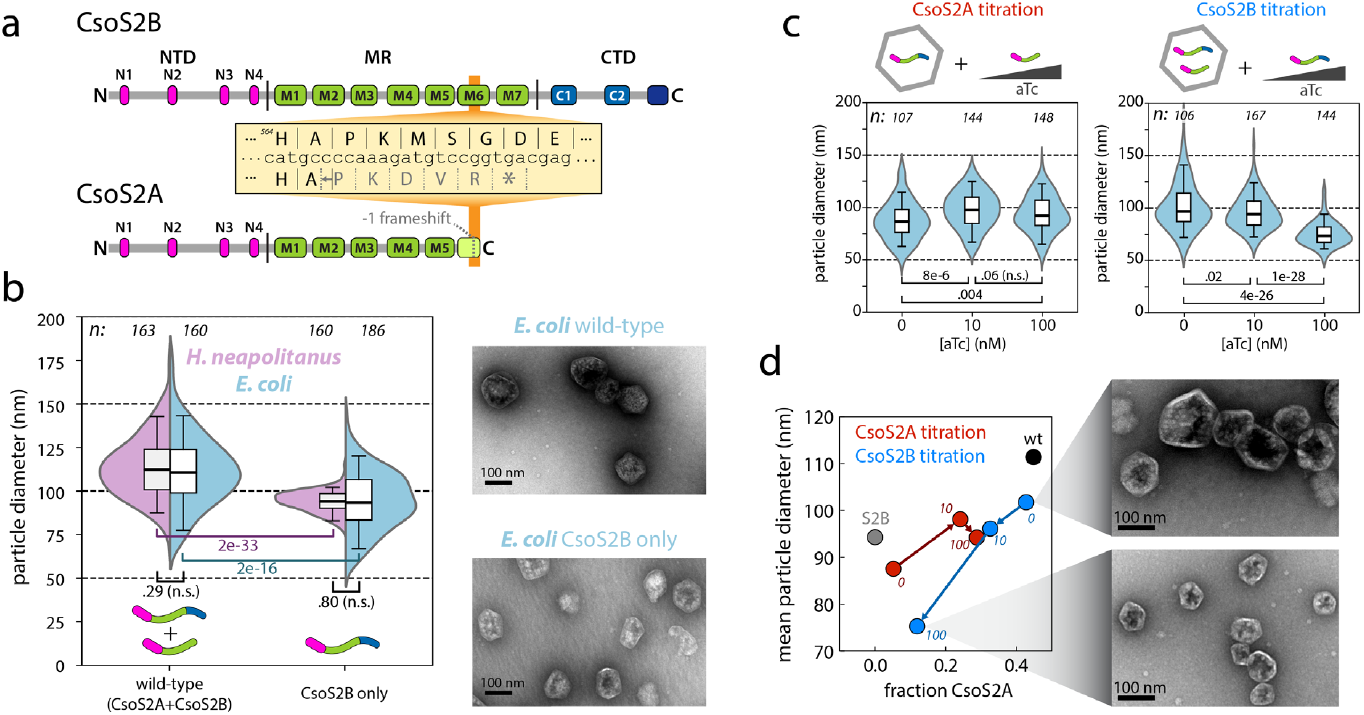
**a**) Diagram of the programmed ribosomal frameshift (PRF) leading to ribosome slippage to -1 frame and premature termination of CsoS2 to generate the CsoS2A isoform. **B**) Kernel density estimate distributions of carboxysome sizes from *H. neapolitanus* and *E. coli* expressing carboxysomes with wtCsoS2 (CsoS2A + CsoS2B) or only CsoS2B in which the PRF has been eliminated. White boxes span the inner quartiles with the median as the central black line. Whiskers are from the 5^th^ to the 95^th^ percentile. Shown below the distributions are p-values for unpaired two-tailed t-tests. Representative negative stain TEM images are shown for the *E. coli* samples. **C**) Scheme and distributions for titration of CsoS2A on a background of CsoS2B-only carboxysomes (red) and titration of CsoS2B on a background of wtCsoS2 carboxysomes. T-test p-values shown below. **D**) Mean particle diameters for titration conditions plotted versus the measured fraction of CsoS2 as CsoS2A from gel densitometry. aTc concentrations in nanomolar are shown in italic next to each data point. Popouts are representative TEM micrographs of the extremes of the CsoS2B titration showing substantially smaller carboxysomes with overexpression of CsoS2B.

Plasmids were transformed into electrocompetent BW25113 *E. coli* cells. For carboxysome expression, cells were grown at 37°C to mid-log (OD600 0.4-0.6) whereupon the temperature was reduced to 18°C and inducer was added: 1 mM IPTG for all samples and a variable amount of anhydrotetracycline (aTc) for the variable co-expression. Cultures were grown overnight then pelleted and frozen or used directly for carboxysome purification.

*H. neapolitanus* cells were grown at 30°C in DSMZ 68 media. The construction of the strain expressing carboxysomes with only CsoS2B from a genomic neutral site under control of a lac operator is described in ref. ^33^. To express carboxysomes and sustain atmospheric growth, this strain was grown in the presence of 1 mM IPTG.

### Carboxysome purification

Carboxysomes were purified as described previously.^28^ The cells from 1L expression cultures were lysed with 25 mL B-PER bacterial lysis reagent (Thermo Fisher) supplemented with 0.01 mg/mL DnaseI, 0.1 mg/mL lysozyme, 1 mM phenylmethylsulfonyl fluoride (PMSF), 10 mM MgCl_2_, and 20 mM NaHCO_3_. Lysate was clarified by centrifugation at 12,000 rcf for 30 min. Carboxysomes were pelleted from the supernatant by centrifugation at 40,000 rcf for 30 min. Crude pelleted carboxysomes were resuspended with 20 mL TEMB buffer (10 mM Tris, 10 mM MgCl_2_, 1 mM EDTA, and 20 mM NaHCO_3_, pH 8.4) on ice and pelleted again at 40,000 rcf for 30 min. The pellet was again gently resuspended in 2mL TEMB and loaded on top of sucrose gradient with 5 mL layers of 10, 20, 30, 40, and 50% sucrose (w/v) in TEMB. The gradient samples were spun in an ultracentrifuge for 35 min at 105,000 rcf. Then the gradients were fractionated into 1 mL fractions and analyzed by SDS-PAGE to determine the carboxysome-containing fractions which also display bluish light scattering from the particles. Fractions with carboxysomes were pooled, centrifuged for 90 min at 105,000 rcf, resuspended with TEMB, and stored at 4°C prior to negative stain transmission electron microscopy (TEM).

### Transmission electron microscopy and size characterization

Carboxysome samples were diluted to a 280 nm absorbance of 0.05-0.1. Prior to sample application, Formvar/carbon coated, copper EM grids were prepared by glow discharge. 5 μL of sample was loaded on to each grid, allowed to sit for at least 2 min, washed, stained with continuous application of 10 uL of 1% (w/v) uranyl acetate solution, and blotted with filter paper. Exposure of the carboxysomes on the grid to air-water interfaces was minimized to prevent carboxysome breakage and increase the accuracy of size analysis. All samples were imaged either with a FEI Tecnai 12 120 kV or JEOL 1200 EX 80 kV transmission electron microscope.

Particle size characterization was performed in ImageJ. Carboxysomes were manually located to avoid particle misidentification and exclude significantly damaged carboxysomes. The images were first given a pixel offset of +4 counts to eliminate any zero count pixels. Then a polyhedron was drawn around each carboxysome and the pixel counts were zeroed within this region. Finally, the particles were selected with a zero count threshold and measured with the automatic particle characterization utility. The minimum Feret diameter–i.e., the closest approach of two parallel lines in contact with the particle projection–was used for all size statistics.

### MR localization to carboxysomes

A series of plasmids expressing MR truncations fused to Superfolder GFP^34^ were each co-transformed with the full carboxysome plasmid (pHnCB10). Carboxysome expression was induced with 1 mM IPTG and GFP fusion expression was induced with 100 nM aTc when the cells reached mid-log phase (OD600 0.4-0.6). The temperature was reduced to 18°C and the cells were grown overnight. Carboxysomes were purified as described above. Then for each sample the absorbance was measured at 340 nm and GFP fluorescence measured with 485 nm excitation and 515 nm emission using a Tecan M1000 plate reader. Due to their size, carboxysomes strongly scatter at 340 nm while soluble protein has low absorption at this wavelength. Thus, the GFP fluorescence intensity divided by the 340 nm absorbance was used as a proxy for the relative amount of GFP loading per carboxysome.

### *Modified CsoS2 in* H. neapolitanus

*H. neapolitanus* is naturally competent and will take up foreign DNA and integrate it into its genome by homologous recombination, provided suitable flanking homology arms. We used this strategy to knock out CsoS2 at the native locus, replacing it with a spectinomycin resistance cassette and validating by PCR. This intermediate strain is incapable of atmospheric growth due to its inability to form carboxysomes. We then knocked in a series of CsoS2 variants having variable M-peptide repeat numbers into the genomic neutral site NS1 along with a kanamycin resistance gene.^33^ All variants could grow in CO_2_-replete conditions with 5% CO_2_. Colony forming unit (CFU) counts were performed by serial dilution of liquid cultures grown at 5% CO_2_ and plated on DSMZ 68 agar plates with spectinomycin (20 μg/mL), kanamycin (10 μg/mL), and IPTG (1 mM) and placed in the corresponding CO_2_ environments: atmosphere (∼0.04%), 0.5%, and 5%.

### Structural modeling of M-peptide shell interaction

The AlphaFold-predicted structures of four divergent CsoS2 proteins were downloaded from Uniprot (RefIDs):^35^ *Halothiobacillus neapolitanus* (O85041), *Prochlorococcus marinus* (Q7V2C8), *Nitrosomonas eutropha* (Q0AHV9), and *Imhoffiella purpurea* (W9VGM3). The coordinates matching the M-peptides were extracted and compared pairwise for structural similarity using TMAlign.^36^ They were commonly aligned to M4 from *N. eutropha* as the nearest structure to the centroid.

*H. neapolitanus* M1-peptide was docked to the luminal side of the junction of three CsoS1A hexamers. The coordinates of this shell trimer were obtained from symmetry expansion of the coordinates of PDB ID 2EWH.^37^ Rosetta was used to independently relax the structures of the M1-peptide and the shell trimer and the best scoring structures were used for protein-protein docking. The M1-peptide was positioned above the three-fold shell axis and roughly aligned with the M1-peptide pseudo-threefold axis. Next the Rosetta local docking routine was used with initial translation randomization of 5Å and orientational randomization of 12°. 500 docked structures were generated.^38^ The best scoring among these clearly fell into a common class of configurations with the VTGs wedged near the symmetry related CsoS1A-His79 residues. The protein-protein interface was further relaxed in Rosetta subject to an applied harmonic distance constraint bringing the VTG threonines into contact with the three-fold His79 residues (Fig. 6c).

### Bioinformatic characterization of M- and C-peptide repeats

All sequences from the Integrated Microbial Genomes database (IMG) with the CsoS2M pfam (PF12288) were downloaded in May 2020 for a total of 770 sequences. These were filtered to eliminate those with partial sequences and unspecified residues. Next the sequences were dereplicated to 95% identity using usearch for a total of 272 unique sequences.^39^ MEME was used to identify peptide motifs and finding, as previously reported, the main known motifs of the N-peptide, M-peptide, C-peptide, and CTP.^40^ MAST from the MEME suite was used to extract the positions of all matches to the M- and C-peptide motifs and the amino acid sequences were extracted for each of these with a 15aa buffer zone on either side. These peptide sequences were aligned all together (i.e. both M- and C-peptides) using mafft.^41^ FastTree was used to generate a phylogenetic tree on the basis of this alignment.^42^ From the tree there are clearly distinguishable clades for the M- and C-peptide classes but some of the initial MAST hits were mislabeled (including *H. neapolitanus* M7). All sequences belonging to each clade were aligned against each other and Weblogo3 was used to generate sequence logos (Fig. 6a, Fig. S1).^43^

## Results

Size in self-assembling biological systems can be controlled by a number of mechanisms including intrinsic curvature built into the geometry of the constituent proteins,^44^ molecular rulers,^45^ excluded volume effects,^18^ and non-equilibrium race conditions between growth and termination.^16^ We hypothesized whether rubisco, which is approximately 65% of the particle by mass and most of the cargo,^5^ might play a pivotal role in guiding the carboxysome size and overall morphology. However, several studies have shown that expression of carboxysome particles without rubisco yields the same basic icosahedral architecture and size, including even in the native *Halothiobacillus* host.^20,46–48^ We repeated this experiment – by recombinantly expressing all carboxysome proteins except rubisco in our *E. coli* heterologous expression system – and obtained similar results (Fig. 2a). In particular, the particle size does not collapse to that of the much smaller complexes formed from shell proteins alone (Fig. 2a, blue lines).

We next turned to the major shell components: the hexameric proteins. An alternative hypothesis is that increasing shell protein concentration would alter the balance between cargo and shell growth in favor of the shell to enclose smaller particles, ultimately short-circuiting the assembly of full-sized particles.^16^ To test this idea, the three major hexameric proteins CsoS1ABC were co-expressed with variable induction alongside the full set of carboxysome proteins. The size and morphologies of the resulting particles were not appreciably different; however, the presence of extra shell proteins was deleterious to the yield of purified carboxysomes (Fig 2b, lysate gel Fig. S2), likely by diverting other carboxysome components into off-pathway assemblies and/or aggregates.

A third hypothesis is that CsoS2, also a major component at ∼15% by mass,^5^ is a possible governor of carboxysome size. To this point, it has previously been found that disordered scaffold proteins can play a role in the size specification of certain icosahedral virus capsids.^49,50^ A particularly striking case is PBCV-1 in which a disordered protein acts as a molecular ruler, effectively measuring the distance between vertices.^45^ We thus tested whether CsoS2 acts as a molecular ruler by inserting a sequence encoding a polyproline II helix – a rigid structural element – of approximately 12 nm between the M3 and M4 repeats.^51^ This construct also produced carboxysomes, but this structural modification of CsoS2 showed no discernable size effect (Fig. 2c).

Given these negative results, we posited that other known biochemistry within the carboxysome could play a role in size control. In *H. neapolitanus* carboxysomes, CsoS2 appears as two distinct isoforms: a short form of ∼61 kDa called CsoS2A and a long form of ∼92 kDa called CsoS2B.^27^ We previously showed that the short form is produced as a result of a -1 programmed ribosomal frameshift (PRF) that leads to the premature termination of CsoS2 (Fig. 3a).^28^ In native *H. neapolitanus* carboxysomes and ones produced recombinantly in *E. coli*, CsoS2A and CsoS2B are present in approximately equimolar quantities. Furthermore, CsoS2B alone can still sustain atmospheric growth in *H. neapolitanus*, while CsoS2A alone cannot due to its inability to assemble purifiable carboxysomes.^28,33^ We thus asked whether the ratio between CsoS2 isoforms has consequences for particle size. Carboxysomes from *H. neapolitanus* were purified from the wild-type strain (having both CsoS2A and B) and a mutant strain in which CsoS2 was knocked out at the native locus and complemented with a CsoS2B-only sequence lacking the frameshifting slippery sequence. Similarly, carboxysomes were expressed and purified from *E. coli* with either the wild-type (wtCsoS2) sequence or the CsoS2B-only sequence in pHnCB10. There was good agreement between the average diameters of carboxysomes obtained from *H. neapolitanus* and *E. coli*. However, a reduction of size was observed in both going from wtCsoS2 to CsoS2B carboxysomes with mean diameters changing from 114 nm to 93 nm for *H. neapolitanus* and 113 nm to 93 nm for *E. coli* (Fig. 3b), similar to the effect of eliminating rubisco (Fig. 2a). While appearing modest, this represents a nearly 40% decrease of available cargo volume in carboxysomes lacking the CsoS2 frameshifting element.

To further explore the connection between CsoS2 isoforms and size, we employed a titratable co-expression system in *E. coli*. In the first, CsoS2A was titrated against a background of carboxysomes with CsoS2B only. In the second, CsoS2B was titrated against a background of wild-type carboxysomes containing both CsoS2A and CsoS2B (Fig. 3cd, lysate gel Fig. S3). Broadly, the results indicate that the balance of CsoS2A and CsoS2B systematically alters the size of the resulting carboxysomes. That is, more CsoS2A correlates with larger particle sizes. Interestingly, overexpression of CsoS2B enhanced the recovered yield of carboxysomes while also reducing the mean particle sizes even lower than that of carboxysomes having only CsoS2B (Fig. 3b, Fig. 3d). This suggests that whatever interaction is driving the size effect may not be fully saturated under normal assembly conditions.

CsoS2A and CsoS2B are distinguished by the absence or presence of the CTD, respectively, which we know to be an indispensable feature for carboxysome assembly.^28^ Wild-type and CsoS2B-only carboxysomes then differ in the relative amount of MR with respect to the CTD. To investigate this effect, we therefore built a series of CsoS2 variants in which we systematically truncated or augmented the MR to include different numbers of the conserved M-peptide repeats. In all of these, we eliminated the PRF sequence such that resulting proteins would be precisely defined with respect to the M-peptide-to-CTD ratio without the complicating effects of frameshifting.

The results showed that each CsoS2 variant produced purifiable carboxysomes that encapsulated rubisco, and, across the series of M-peptide numbers, we observed a dramatic size trend (Fig. 4, pooled carboxysome gel Fig. S4). The smallest carboxysomes were those with no M-peptide repeats and with an average diameter of just over 40 nm, a size which is highly similar to the single particle structure solved by Ni and Jiang et al.^21^ at T=9 and 36.9 nm diameter. In contrast, augmenting CsoS2B with copies of M-peptides M1-M6 resulted in particles having substantially larger diameters, with an average of 130 nm versus 94 nm for unmodified CsoS2B.

**Figure 4.**
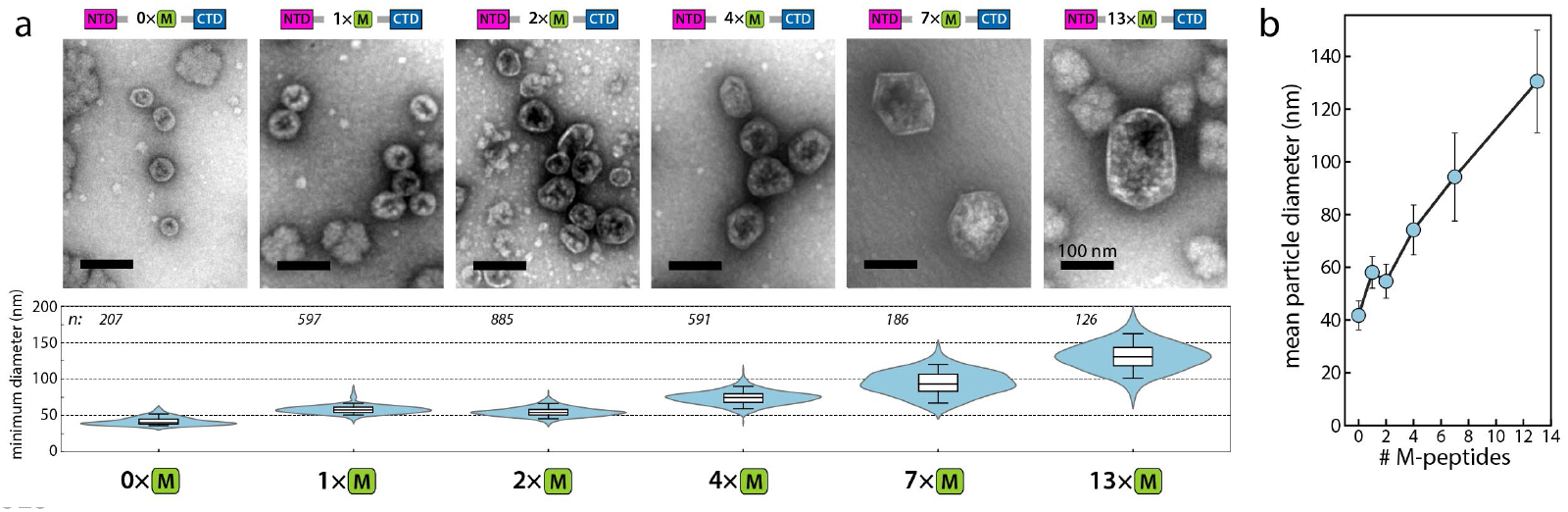
**a**) A series of carboxysomes were produced with variable numbers of M-peptide repeats going from 0 to 13 (CsoS2B has 7). Negative stain TEM images are shown for representative particles in the population. All images are at the same scale with 100 nm scale bars. Below is the distribution of minimum particle Feret diameters. Central black line indicates the median, white box represents inner quartiles, and whiskers extend to the 5^th^ to 95^th^ percentile. **B**) Plot of mean particle diameters as a function of the number of M-peptides. Whiskers are +/-one standard deviation of the distributions.

In addition, we tested a subset of these variants for complementation in the wtCsoS2 knockout *H. neapolitanus* strain. While all strains could grow under 5% CO_2,_ we found that the MR truncations negatively impacted survival at atmospheric CO_2_ concentrations (∼0.04%) with 1 and 4 M-peptide repeats unable to sustain growth while 7 and 13 M-peptide repeats could. At 0.5% CO_2_ all but the 1 M-peptide strain could survive (Fig. S5).

The two C-peptides of the CTD are related to the M-peptides and share a common motif of three VTG triplets each separated by 8 residues (Fig. S1). This similarity leads us to ask – do these C-peptides play a similar role as the M-peptides with respect to carboxysome size? To test this hypothesis, we created two CsoS2 variants: one with a duplication of both C-peptide repeats and another with duplication of the entire CTD, including the CTP. In contrast to the MR augmentation which produced substantially larger carboxysomes, the C-peptide duplications did not result in larger particles and, instead, produced similarly sized or even slightly smaller carboxysomes compared to CsoS2B (Fig. S6).

It is now well established that the CTD forms interactions with the carboxysome shell via both the C-peptides and CTP.^20,21^ Our recent study found that the MR also specifically interacts with the shell hexamers and that the VTG triplets are an essential component of that interaction.^29^ To further test this interaction in the context of carboxysome assembly we designed a series of MR-GFP fusion proteins having different numbers of M-peptides. These were co-expressed alongside complete carboxysomes, and we quantified the relative GFP loading (Fig. S7). We found a strong dependence on the M-peptide repeat number, with more repeats resulting in more loading. The quantity of any of these fusions in the carboxysomes as visualized by SDS-PAGE was far below that of CsoS2 such that they did not appreciably alter the balance of M- and C-peptides. Together, these results imply that the MR interacts with the shell during assembly and does so in a multivalent fashion.

In summary, the M-peptides and C-peptides both share similar sequence features and interact with the shell. However, they have divergent effects on the size: the number of M-peptides strongly dictates the particle size while the C-peptides are absolutely required for carboxysome-like particle formation, but their relative numbers seemingly do not alter particle size.

## Discussion

In this study we have examined the factors determining the size of α-carboxysomes. We discovered that the particle size is strongly specified by the M-peptide repeat number, and that this size preference is transmitted via its interaction with the shell. Below, we discuss the implications of these observations and put forward a size control model with a potential structural mechanism.

The dispensability of rubisco to the formation of carboxysome-like particles implies that it is not an active participant in the assembly itself. Instead, it acts more as a client protein, partitioning into a phase-separated condensate with CsoS2 via the NTD on the surface of the growing shell.^14^ Thus we are left with the interplay between CsoS2 and shell from which to build a model. This model must take into account several observations: 1) the CTD and MR both interact with shell proteins;^20,21,29^ 2) their interactions are both multivalent in nature and due to the repeated M- and C-peptides, respectively^21,29^ (Fig. S7), 3) the M- and C-peptides share a sequence motif – the [V/I]TG triplet – so that they likely have related modes of contacting the shell (Fig. S1); and 4) the M- and C-peptides differ in their effects on particle size, i.e. M-peptides encourage larger particles while C-peptides are size neutral (Fig. 4, Fig. S6). Taken together, we believe that these factors point towards a model in which CsoS2 modifies the mechanical properties of the shell.

It has been previously shown that expressing shell proteins alone results in small icosahedral structures, revealing that, in the absence of other factors, the capsomers form stable high-curvature contacts.^19,52,53^ Introduction of CsoS2 into this mix frustrates the formation of these small particles and instead produces significantly larger particles with lower curvature at the shell-shell interfaces. We specifically propose that the essence of the size control lies in energetic differences of shell binding between the M- and C-peptides with respect to the angles at the junctions of shell hexamers. This energetic effect can be expressed as two related but distinct models we call “preferred curvature” and “flexible curvature” (Fig. 5a). In preferred curvature, the C-peptide has an energy minimum at a greater shell contact angle than the M-peptide. In the flexible curvature model the C-peptide and M-peptide have the same energy minimum angle, however, the C-peptide has a broader energy function and can tolerate greater contact angles than the M-peptide. Note that the angle, while represented as a two-dimensional plane angle in the cartoon schematic (Fig. 5a), is likely a three-dimensional solid angle. Finally, it should be emphasized that the model is not for C-peptide binding at shell junctions directly bordering the pentameric vertex capsomer – where Ni and Jiang et al. find the CTP^21^ – but rather for stabilizing hexamer-only junctions in the vicinity of the vertex that must sustain higher curvature than those further away for which the M-peptide is favored (Fig. 5b).^54^ It is possible that the CTP interaction plays a supporting role in the size control, however, it is likely minor since carboxysomes made without pentameric capsomers form with typical minimum diameters.^6^

**Figure 5.**
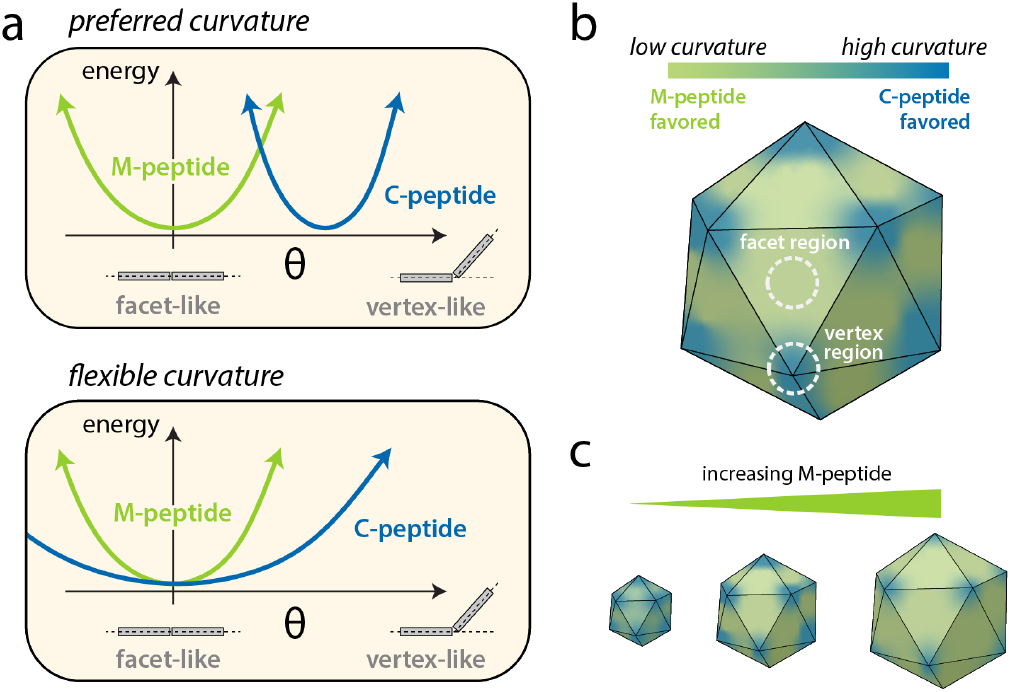
**a**) Energetic models of M- and C-peptide shell interactions. Plots show potential energy of shell-bound M- (green) and C-peptides (blue) as a function of shell angle. In the “preferred curvature” model the C-peptide has an energy minimum for a high curvature shell configuration. In the “flexible curvature” model the C-peptide has the same preferred angle as the M-peptide but a shallower energy well. **b**) Idealized map of carboxysome shell curvature. Higher curvature regions are centered on vertices while lower curvature prevails in between and on facets. The model predicts green regions will be more favorable for M-peptide binding and blue regions will be more favorable for C-peptide binding. **c**) More M-peptide stabilizes particles with a greater share of low curvature surface and thus greater diameter.

This model can be used to interpret a number of experimental observations. First, the M-peptide series in Fig. 4 entails increasing M-peptide numbers on a constant background of two C-peptides. To a first approximation, the number of high curvature sites will be roughly the same as particles increase in size while the number of low curvature sites will scale with the surface area. Thus, smaller particles are energetically better suited to a low M- to C-peptide ratio, while larger particles will benefit from a high M- to C-peptide ratio (Fig. 5c).^54^ Second, the insufficiency of CsoS2A alone to form carboxysomes may be due to the fact that, while it could perhaps knit together flat sheets of hexamer facets with M-peptides, it does not possess the C-peptides necessary to stabilize the regions of highest curvature near the vertices and is thus incapable of producing closed particles. Finally, the larger size of *H. neapolitanus* carboxysomes with wtCsoS2 is explained by the extra abundance of the M-peptides relative to CsoS2B-only to extend the low curvature between vertices.

At first glance it appears improbable that one could make structural inferences on the basis of our phenomenological model given CsoS2’s high disorder score throughout the sequence.^13,14,28^ However, analysis of diverse CsoS2 sequence structural predictions from the AlphaFold Uniprot database revealed that, while the overall structures are amorphous and low confidence, the aligned M-peptide sequences give a remarkably consistent microdomain (Fig. 6a). This structure has the VTG triplet motifs arranged into a triangular structure with pseudo-threefold symmetry. Furthermore, the conserved pair of cysteine residues are located in close proximity to one another. This is notable because the carboxysome interior is thought to possess an oxidized environment, and thus one might expect a fold-stabilizing disulfide bridge to form.^9,55,56^

**Figure 6.**
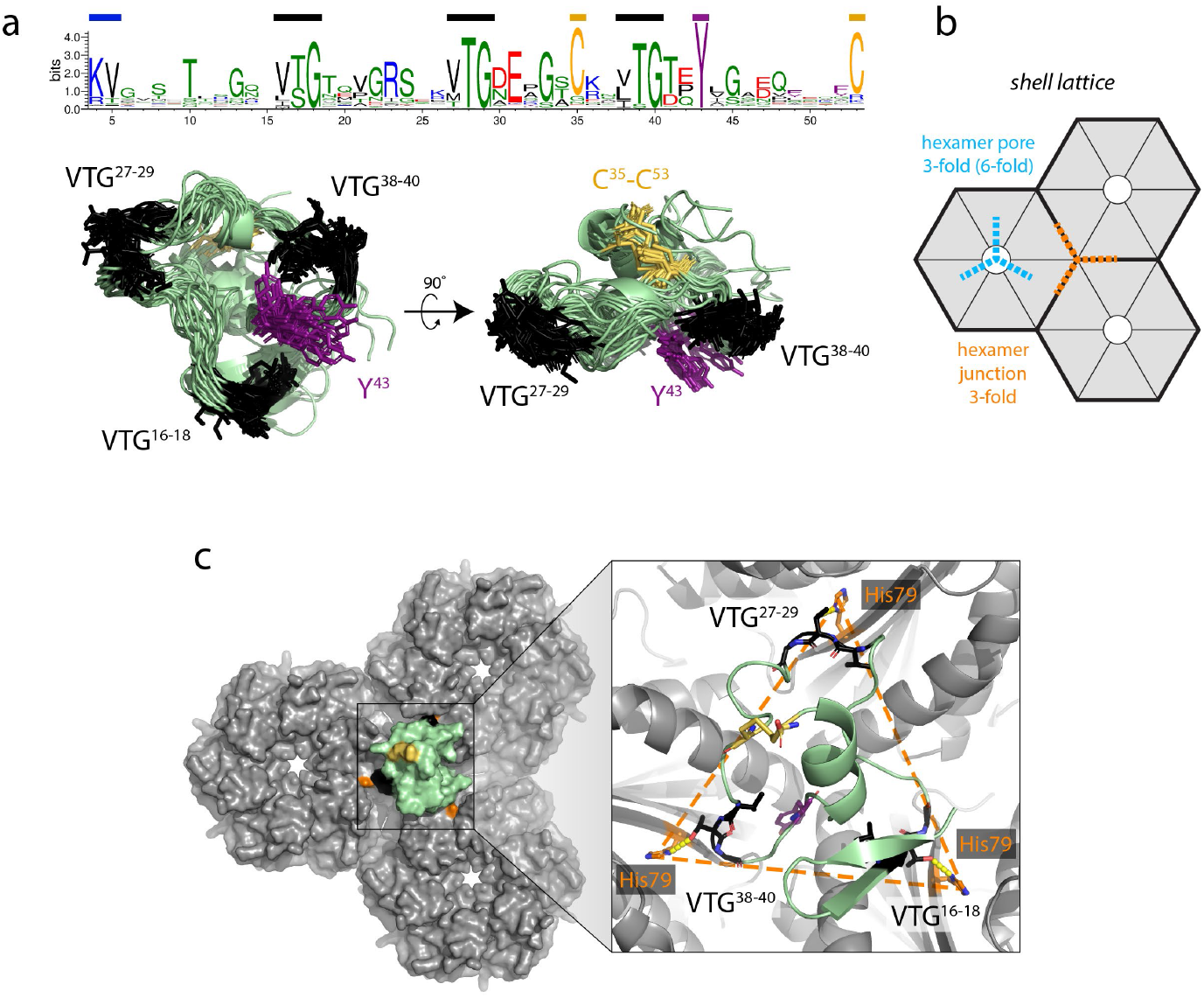
**a**) Sequence logo for M-peptide created from 1662 M-peptide sequences pulled from 272 CsoS2 sequences after dereplication to 95% identity. Aligned AlphaFold predictions for all M-peptides in CsoS2s from *Halothiobacillus neapolitanus, Prochlorococcus marinus, Nitrosomonas eutropha*, and *Imhoffiella purpurea*. Key conserved residues are colored matching the bars on the sequence logo. **b**) Schematic of hexameric shell lattice indicating the site classes with three-fold symmetry. **c**) Surface representation of *H. neapolitanus* M1-peptide docked onto the three-fold hexamer junction with Rosetta. Zoom in shows the VTG motifs in contact with the triangle defined by CsoS1A-His79.

The above structural prediction combined with the curvature model suggests that this pseudo-threefold symmetric M-peptide ought to bind at a shell junction with three-fold symmetry (Fig. 6b, orange). We took the AlphaFold prediction for the *H. neapolitanus* CsoS2 M1-peptide and used Rosetta to dock the structure and relax the interface onto the three-fold hexamer junction using the CsoS1A coordinates (PDB ID: 2EWH).^37,38^ The fit has good size and shape complementarity with the depression at the junction. A cryoEM structure with portions of the CTD resolved was very recently described by Ni and Jiang et al.^21^ Most interestingly, they identified CsoS1A-His79 as an important contact for CTD binding through the VTG motifs. His79 lies in close proximity to the triad of VTGs of the docked M1-peptide. Using harmonic distance constraints in Rosetta the M1-peptide was relaxed into an idealized geometry with the VTG threonine hydroxyls each in hydrogen bonding contact with the His79 delta nitrogen while maintaining the same basic tertiary fold (Fig. 6c).

What might structurally distinguish the C-peptide from the M-peptide and explain their energetic preferences for different shell curvatures? The M-peptide consensus sequence is essentially the C-peptide consensus plus several additional conserved positions, namely the aforementioned cysteine pair, a tyrosine residue appearing three residues downstream of the last VTG motif, and a lysine-valine pair near the beginning (Fig. 6a, Fig. S1). In a recent study of the MR, we observed significant effects of the conserved tyrosine on the biochemical interactions with the shell and sufficiency of resulting carboxysomes for atmospheric growth in *H. neapolitanus*, highlighting a distinct yet crucial role of the M-peptides.^29^ The AlphaFold predictions for the C-peptides are of low confidence and lack a common structure. The CTD-shell structure from Ni and Jiang et al. shows that even as the C-peptide and CTP residues are well resolved, particularly at the VTG-binding contacts, the overall structure threads its way along the hexamers without a clear tertiary structure of its own.^21^ We hypothesize, in contrast, that the M-peptide acts more as a microdomain with a characteristic fold (Fig. 6a). Taken together, this suggests the M-peptide motif is more rigid and prefers to bind low-angle hexamer junctions while the C-peptide motif is more flexible and can accommodate higher angle junctions.

Given this model of size we can make some inferences about the natural diversity of CsoS2 sequences and the attending size consequences. Broadly speaking, we predict that organisms with high M- to C-peptide ratios will produce on average larger carboxysomes while those with low M- to C-peptide ratios will produce smaller carboxysomes. Furthermore, we also predict that, all else being equal, CsoS2 sequences containing a PRF will afford larger carboxysomes than an otherwise identical sequence. It will be interesting to see whether these predictions map onto environmental conditions. For example, we observed that CsoS2 truncated to 4 M-peptides could not sustain growth in *H. neapolitanus* under atmospheric CO_2_ concentrations but could at 0.5% CO_2_ (Fig. S5).

More broadly, how does this mechanism of size control compare to other bacterial microcompartments (BMCs)? β-carboxysomes do not form large icosahedral shell structures in the absence of rubisco^52^ nor do they possess an obvious analogue to CsoS2. Combined with the evidence for inside-out assembly, this suggests that the size is determined as a result of a close interplay between a growing shell stabilized around a dense cargo.^9,16,17^ Among other BMCs (e.g., Pdu and Eut) there is a wide variety of sizes and morphologies. It is not presently clear whether they fall under common assembly principles and size controls. Interestingly, some BMC shell proteins can assemble into diverse macrostructures such as nanotubes, extended sheets, and rosettes.^57^ A leading hypothesis for the BMC size control is that it may be driven by the relative stoichiometries of constituent shell capsomers that differ in their toleration of curvature.^19,54,58,59^ With additional structural and biochemical data, it will thus be interesting to see whether there is a unifying principle for size control of microcompartments broadly or if, as with viruses, there are multiple mechanistic routes to achieve similar outcomes.

## Conclusions

In this study we sought to uncover the determinants of particle size in α-carboxysomes. We discovered that the size is set primarily by the scaffold protein CsoS2 and its interactions with the shell. Through systematic modification of CsoS2 we observed that the size effect is mediated by the relative numbers of M-peptide repeats found in the MR and the C-peptide repeats found in the CTD. Additionally, we determined that a functional consequence of the PRF in *H. neapolitanus* CsoS2 is to increase particle size at no cost of additional sequence.

We propose an energetic model based on differential curvature preferences between the M- and C-peptides and connect this model to a mechanistic structural hypothesis on the basis of AlphaFold structural predictions and new experimental breakthroughs in carboxysome structure.

An increasingly comprehensive view of α-carboxysome assembly is coming into focus as the structures and interactions linking the components together are elucidated. Only a few key gaps – most notably the CsoS2-MR shell interaction – remain before a complete account of the structure, spanning atomic detail to functional particles. With this depth of information we think that the carboxysome is a useful paradigm for complex self-assembly and potentially a template, physically but also conceptually, for future engineering efforts.

## Acknowledgements

We thank Dr. Hanlun Jiang for assistance with Rosetta modeling. We acknowledge the staff at the UC Berkeley Electron Microscope Laboratory for training and assistance with TEM. This work was supported by the U. S. Department of Energy, Office of Science, Office of Basic Energy Sciences, Chemical Sciences, Geosciences, & Biosciences Division, Physical Biosciences Program, under Award Number DE-SC0016240 to D.F.S. and D.F.S. is an Investigator of the Howard Hughes Medical Institute.

## Footnotes

§ These studies with shell-only complexes use a subset of capsomer paralogs. It is unknown whether other combinations might change size characteristics.

