## Supplementary Information for "α-carboxysome size is controlled by the disordered scaffold protein CsoS2"

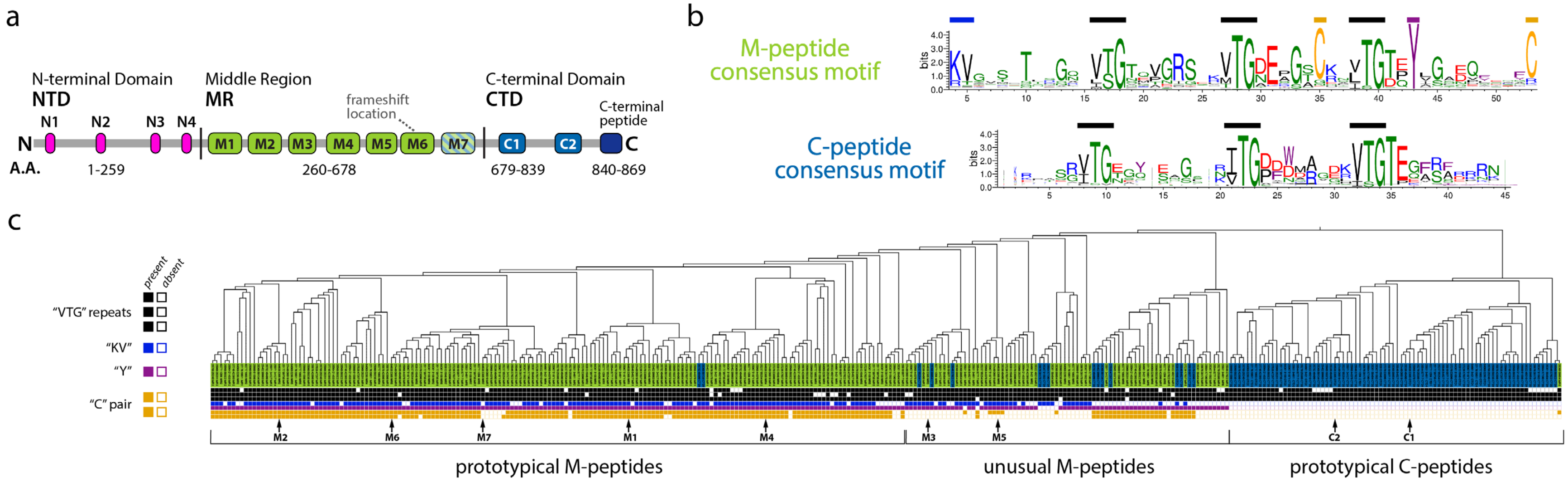

***Figure S1***

**a**) CsoS2 has been divided into three domains based on the self-similarity of the repeated peptides contained within each. We suggest an update to the repeat nomenclature in which M7, known previously as R7, fits better as an M-peptide rather than a C-peptide. We aligned all M- and C-peptide sequences located using the motifs generated by MEME and built a phylogenetic tree (c). (Note that the tree is heavily pruned from the full set of sequences in the alignment which has a total of 2190 members.) In this representation M7 clearly belongs with the other M-peptide repeats while C1 and C2 (previously known as R8 and R9) are definitively in the distinct C-peptide clade. The colors of the leaves are according to the original identification by MAST. **b**) Sequence logos created with Weblogo3 for the M-peptide and C-peptide sequences based on clade membership. These clearly show the common element of both sequences as the VTG triplet. The M-peptide has additional conserved features. **c**) Phylogenetic tree of M- and C-peptide repeats visualized with iTOL. We highlight the presence or absence of each of the conserved features across the variety of repeats and this makes a further subdivision stand out in the M-peptides between those that carry most of the conserved features and those missing a subset.

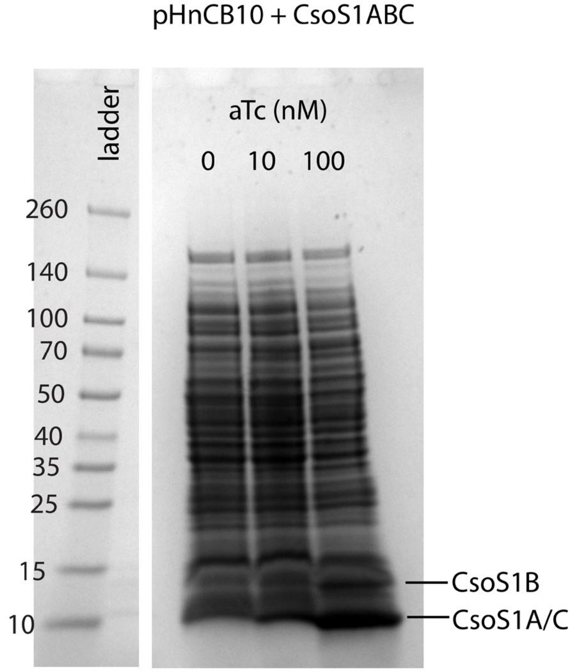

***Figure S2***

SDS-PAGE analysis of shell titration raw cell lysates from main text Fig. 2b. Shell hexamers (CsoS1ABC) are expressed with variable anhydrotetracycline (aTc) inducer concentrations alongside full carboxysome expression (pHnCB10).

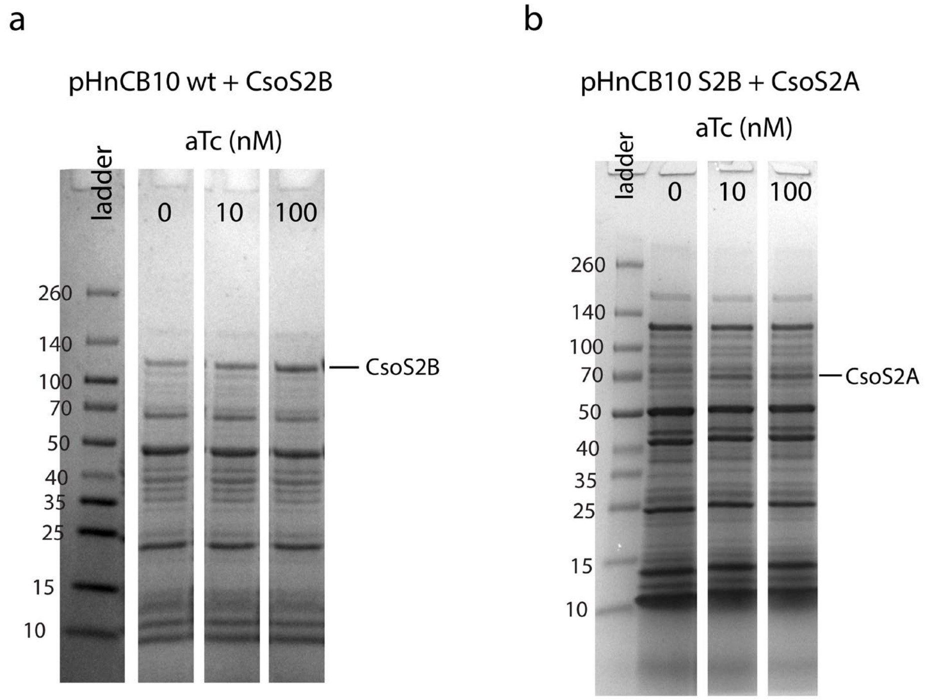

***Figure S3***

SDS-PAGE analysis of CsoS2 isoform titration raw cell lysates from main text Fig. 3c. a) CsoS2B was titrated with variable aTc concentrations alongside wild-type carboxysomes (pHnCB10 wt). B) CsoS2A was titrated with variable aTc concentrations alongside carboxysomes with only the CsoS2B isoform (pHnCB10 S2B).

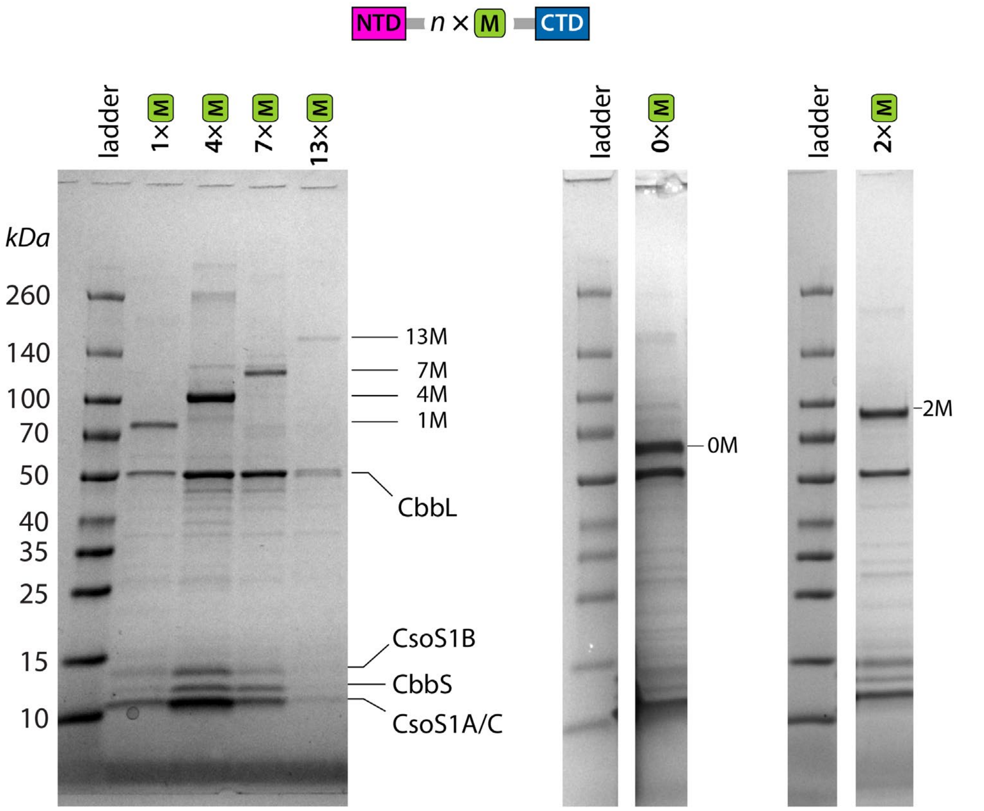

***Figure S4***

SDS-PAGE gels of pooled purified carboxysome samples for the variable M-peptide number size series from main text Fig. 4.

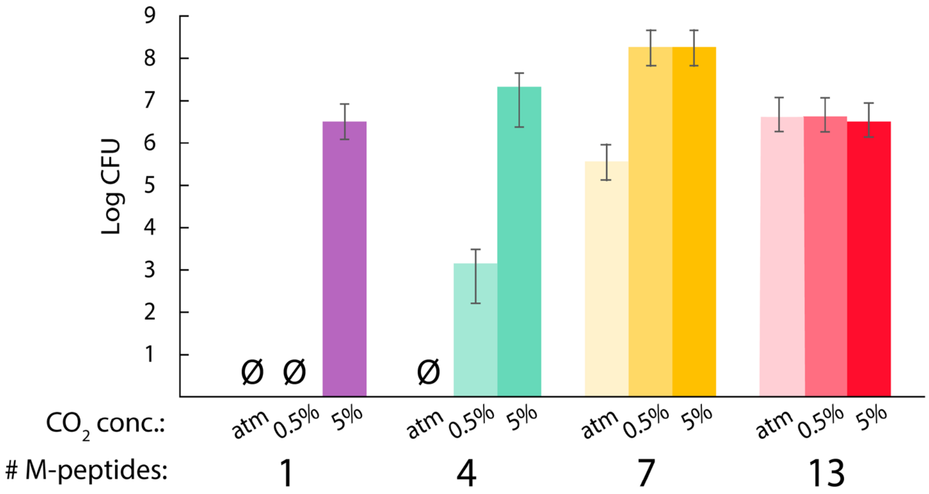

***Figure S5***

Growth of *H. neapolitanus* with CsoS2 variants containing different numbers of M-peptide repeats. CFU counts are the geometric means for triplicates of the plating assay. Error bars represent +/- one geometric standard deviation.

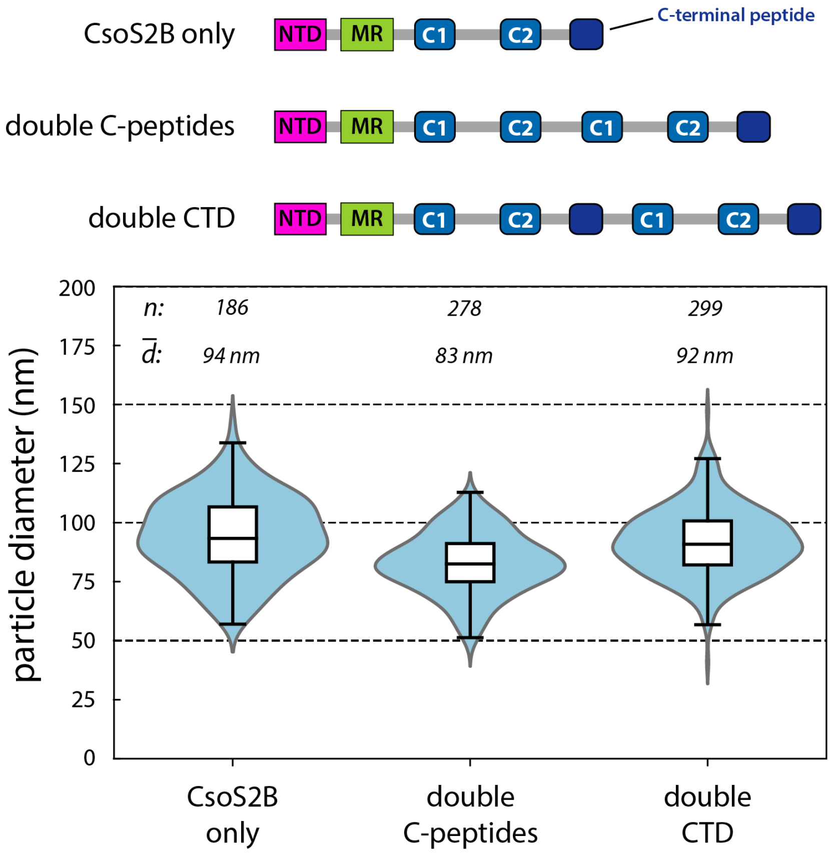

***Figure S6***

Size effects of CTD augmentation. Three CsoS2 variants were compared to examine the effect of increasing the C-peptide repeat count: CsoS2B, double C-peptides with duplication only of the C-peptides, and double CTD with duplication of the entire CTD including the CTP. Kernel density estimations of the particle diameter distributions are shown in light blue with the central black line at the median and the white box spanning the inner quartiles. Whiskers are from the 5th to 95th percentile. Particle numbers and mean diameters are indicated along the top.

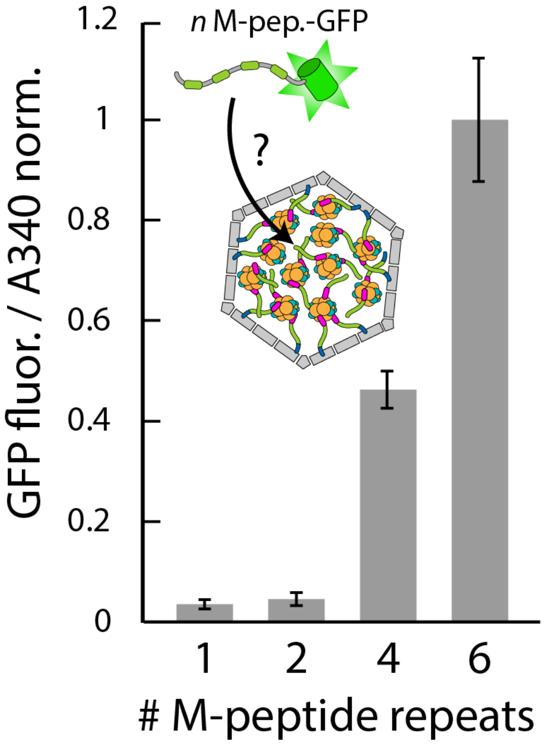

***Figure S7***

Targeting assay for localization of MR-GFP fusions with different numbers of M-peptide repeats. The fluorescence is proportional to the amount of GFP and the absorbance at 340 nm is due primarily to the scattering of carboxysomes and thus the ratio is an estimate of the loading efficiency. Error bars are +/- one standard deviation of three independent carboxysome preparations.

***Table S1***

Plasmids used for protein expression.

| ID | description | back-bone | anti-biotic | CsoS2 or CsoS2 fragment sequence |
|---|---|---|---|---|
| Lz102 | full carboxysome with wtCsoS2 | pHnCB | Cm | MPSQSGMNPADLSGLSGKELARARRAALSKQGKAAVSNKTASVNRSTKQAASSINTNQV<br>RSSVNEVPTDYQMADQLCSTIDHADFGTESNRVRDLCRQRREALSTIGKKAVKTNGKPSG<br>RVRPQQSVVHNDAMIENAGDTNQSSSTSLNNELSEICSIADDMPERFGSQAKTVRDICRA<br>RRQALSERGTRAVPPKPQSQGGPGRNGYQIDGYLDTALHGRDAAKRHREMLCQYGRGT<br>APSCKPTGRVKNSVQSGNAAPKKVETGHTLSGGSVTGTQVDRKSHVTGNEPGTCRAVT<br>GTEYVGTEQFTSFCNTSPKPNATKVNVTTTARGRPVSGTEVSRTEKVTGNESGVCRNVT<br>GTEYMSNEAHFSLCGTAAKPSQADKVMFGATARTHQVVSGSDEFRPSSVTGNESGAKRT<br>ITGSQYADEGLARLTINGAPAKVARTHTFAGSDVTGTEIGRSTRVTGDESGSCRSISGTEY<br>LSNEQFQSFCDTKPQRSPFKVGQDRTNKGQSVTGNLVDRSELVTGNEPGSCSRVTGSQ<br>YGQSKICGGGVGKVRSMRTLRGTSVSGQQLDHA^PKMSGDERGGCMPVTGNEYYGRE<br>HFEPFCTSTPEPEAQSTEQSLTCEGQIISGTSVDASDLVTGNEIGEQQLISGDAYVGAQQT<br>GCLPTSPRFNQTGNVQSMGFKNTNQPEQNFAPGEVMPTDFSIQTPARSAQNRITGNDIAP<br>SGRITGPGMLATGLITGTPEFRHAARELVGSPQPMAMAMANRNKAAQAPVVQPEVVATQ<br>EKPELVCAPRSDQMDRVSGEGKERCHITGDDWSVNKHITGTAGQWASGRNPSMRGNAR<br>VVETSAFANRNVPKPEKPGSKITGSSGNDTQGSLITYSGGARG*<br>frameshift: ^ (PKDVR*) |
| Lz111 | full carboxysome with CsoS2B only | pHnCB | Cm | MPSQSGMNPADLSGLSGKELARARRAALSKQGKAAVSNKTASVNRSTKQAASSINTNQV<br>RSSVNEVPTDYQMADQLCSTIDHADFGTESNRVRDLCRQRREALSTIGKKAVKTNGKPSG<br>RVRPQQSVVHNDAMIENAGDTNQSSSTSLNNELSEICSIADDMPERFGSQAKTVRDICRA<br>RRQALSERGTRAVPPKPQSQGGPGRNGYQIDGYLDTALHGRDAAKRHREMLCQYGRGT<br>APSCKPTGRVKNSVQSGNAAPKKVETGHTLSGGSVTGTQVDRKSHVTGNEPGTCRAVT<br>GTEYVGTEQFTSFCNTSPKPNATKVNVTTTARGRPVSGTEVSRTEKVTGNESGVCRNVT<br>GTEYMSNEAHFSLCGTAAKPSQADKVMFGATARTHQVVSGSDEFRPSSVTGNESGAKRT<br>ITGSQYADEGLARLTINGAPAKVARTHTFAGSDVTGTEIGRSTRVTGDESGSCRSISGTEY<br>LSNEQFQSFCDTKPQRSPFKVGQDRTNKGQSVTGNLVDRSELVTGNEPGSCSRVTGSQ<br>YGQSKICGGGVGKVRSMRTLRGTSVSGQQLDHAPKMSGDERGGCMPVTGNEYYGREH<br>FEPFCTSTPEPEAQSTEQSLTCEGQIISGTSVDASDLVTGNEIGEQQLISGDAYVGAQQTG<br>CLPTSPRFNQTGNVQSMGFKNTNQPEQNFAPGEVMPTDFSIQTPARSAQNRITGNDIAPS<br>GRITGPGMLATGLITGTPEFRHAARELVGSPQPMAMAMANRNKAAQAPVVQPEVVATQE<br>KPELVCAPRSDQMDRVSGEGKERCHITGDDWSVNKHITGTAGQWASGRNPSMRGNARV<br>VETSAFANRNVPKPEKPGSKITGSSGNDTQGSLITYSGGARG* |

| ID | description | back-bone | anti-biotic | CsoS2 or CsoS2 fragment sequence |
|---|---|---|---|---|
| Az42 | carboxysome without rubisco | pHnCB | Cm | same as Lz102 |
| Az45 | CsoS1ABC for shell titration | pFA | Kan | N/A |
| Az08 | full carboxysome with polyproline CsoS2 | pHnCB | Cm | MPSQSGMNPADLSGLSGKELARARRAALSKQGKAAVSNKTASVNRSTKQAASSINTNQV<br>RSSVNEVPTDYQMADQLCSTIDHADFGTESNRVRDLCRQRREALSTIGKKAVKTNGKPSG<br>RVRPQQSVVHNDAMIENAGDTNQSSSTSLNNELSEICSIADDMPERFGSQAKTVRDICRA<br>RRQALSERGTRAVPPKPQSQGGPGRNGYQIDGYLDTALHGRDAAKRHREMLCQYGRGT<br>APSCKPTGRVKNSVQSGNAAPKKVETGHTLSGGSVTGTQVDRKSHVTGNEPGTCRAVT<br>GTEYVGTEQFTSFCNTSPKPNATKVNVTTTARGRPVSGTEVSRTEKVTGNESGVCRNVT<br>GTEYMSNEAHFSLCGTAAKPSQADKVMFGATARTHQVVSGSDEFRPSSVTGNESGAKRT<br>ITGSQYADEGLARLTSGGPPPPPPPPPPAPAPAPAPPPPPPPPPPAPAPAPAPPPPPPPPPPGG<br>GSTINGAPAKVARTHTFAGSDVTGTEIGRSTRVTGDESGSCRSISGTEYLSNEQFQSFCDT<br>KPQRSPFKVGQDRTNKGQSVTGNLVDRSELVTGNEPGSCSRVTGSQYGQSKICGGGVG<br>KVRSMRTLRGTSVSGQQLDHAPKMSGDERGGCMPVTGNEYYGREHFEPFCTSTPEPEA<br>QSTEQSLTCEGQIISGTSVDASDLVTGNEIGEQQLISGDAYVGAQQTGCLPTSPRFNQTGN<br>VQSMGFKNTNQPEQNFAPGEVMPTDFSIQTPARSAQNRITGNDIAPSGRITGPGMLATGLI<br>TGTPEFRHAARELVGSPQPMAMAMANRNKAAQAPVVQPEVVATQEKPELVCAPRSDQM<br>DRVSGEGKERCHITGDDWSVNKHITGTAGQWASGRNPSMRGNARVVETSAFANRNVPK<br>PEKPGSKITGSSGNDTQGSLITYSGGARG* |
| Lz110 | CsoS2A for titration | pFA | Kan | MPSQSGMNPADLSGLSGKELARARRAALSKQGKAAVSNKTASVNRSTKQAASSINTNQV<br>RSSVNEVPTDYQMADQLCSTIDHADFGTESNRVRDLCRQRREALSTIGKKAVKTNGKPSG<br>RVRPQQSVVHNDAMIENAGDTNQSSSTSLNNELSEICSIADDMPERFGSQAKTVRDICRA<br>RRQALSERGTRAVPPKPQSQGGPGRNGYQIDGYLDTALHGRDAAKRHREMLCQYGRGT<br>APSCKPTGRVKNSVQSGNAAPKKVETGHTLSGGSVTGTQVDRKSHVTGNEPGTCRAVT<br>GTEYVGTEQFTSFCNTSPKPNATKVNVTTTARGRPVSGTEVSRTEKVTGNESGVCRNVT<br>GTEYMSNEAHFSLCGTAAKPSQADKVMFGATARTHQVVSGSDEFRPSSVTGNESGAKRT<br>ITGSQYADEGLARLTINGAPAKVARTHTFAGSDVTGTEIGRSTRVTGDESGSCRSISGTEY<br>LSNEQFQSFCDTKPQRSPFKVGQDRTNKGQSVTGNLVDRSELVTGNEPGSCSRVTGSQ<br>YGQSKICGGGVGKVRSMRTLRGTSVSGQQLDHAPKDVR* |
| Az44 | CsoS2B for | pFA | Kan | same as Lz111 |

| ID | description | back-bone | anti-biotic | CsoS2 or CsoS2 fragment sequence |
|---|---|---|---|---|
| | titration | | | |
| Lz124 | full carboxysome, 0M CsoS2 | pHnCB | Cm | MPSQSGMNPADLSGLSGKELARARRAALSKQGKAAVSNKTASVNRSTKQAASSINTNQV RSSVNEVPTDYQMADQLCSTIDHADFGTESNRVRDLCRQRREALSTIGKKAVKTNGKPSG RVRPQQSVVHNDAMIENAGDTNQSSSTSLNNELSEICSIADDMPERFGSQAKTVRDICRA RRQALSERGTRAVPPKPQSQGGPGRNGYQIDGYLDTALHGRDAAKRHREMLCQYGRGT APSCKPTGRVKNSVQSGNAAPKTSPRFNQTGNVQSMGFKNTNQPEQNFAPGEVMPTDF SIQTPARSAQNRITGNDIAPSGRITGPGMLATGLITGTPEFRHAARELVGSPQPMAMAMAN RNKAAQAPVVQPEVVATQEKPELVCAPRSDQMDRVSGEGKERCHITGDDWSVNKHITGT AGQWASGRNPSMRGNARVVETSAFANRNVPKPEKPGSKITGSSGNDTQGSLITYSGGAR G* |
| Bz125 | full carboxysome, 1M CsoS2 | pHnCB | Cm | MPSQSGMNPADLSGLSGKELARARRAALSKQGKAAVSNKTASVNRSTKQAASSINTNQV RSSVNEVPTDYQMADQLCSTIDHADFGTESNRVRDLCRQRREALSTIGKKAVKTNGKPSG RVRPQQSVVHNDAMIENAGDTNQSSSTSLNNELSEICSIADDMPERFGSQAKTVRDICRA RRQALSERGTRAVPPKPQSQGGPGRNGYQIDGYLDTALHGRDAAKRHREMLCQYGRGT APSCKPTGRVKNSVQSGNAAPKTSTPEPEAQSTEQSLTCEGQIISGTSVDASDLVTGNEIG EQQLISGDAYVGAQQTGCLPTSPRFNQTGNVQSMGFKNTNQPEQNFAPGEVMPTDFSIQ TPARSAQNRITGNDIAPSGRITGPGMLATGLITGTPEFRHAARELVGSPQPMAMAMANRN KAAQAPVVQPEVVATQEKPELVCAPRSDQMDRVSGEGKERCHITGDDWSVNKHITGTAG QWASGRNPSMRGNARVVETSAFANRNVPKPEKPGSKITGSSGNDTQGSLITYSGGARG* |
| Az41 | full carboxysome, 2M CsoS2 | pHnCB | Cm | MGSNMPSQSGMNPADLSGLSGKELARARRAALSKQGKAAVSNKTASVNRSTKQAASSIN TNQVRSSVNEVPTDYQMADQLCSTIDHADFGTESNRVRDLCRQRREALSTIGKKAVKTNG KPSGRVRPQQSVVHNDAMIENAGDTNQSSSTSLNNELSEICSIADDMPERFGSQAKTVRD ICRARRQALSERGTRAVPPKPQSQGGPGRNGYQIDGYLDTALHGRDAAKRHREMLCQY GRGTAPSCKPTGRVKNSVQSGNAAPKKVETGHTLSGGSVTGTQVDRKSHVTGNEPGTC RAVTGTEYVGTEQFTSFCNTSPTSTPEPEAQSTEQSLTCEGQIISGTSVDASDLVTGNEIG EQQLISGDAYVGAQQTGCLPTSPRFNQTGNVQSMGFKNTNQPEQNFAPGEVMPTDFSIQ TPARSAQNRITGNDIAPSGRITGPGMLATGLITGTPEFRHAARELVGSPQPMAMAMANRN KAAQAPVVQPEVVATQEKPELVCAPRSDQMDRVSGEGKERCHITGDDWSVNKHITGTAG QWASGRNPSMRGNARVVETSAFANRNVPKPEKPGSKITGSSGNDTQGSLITYSGGARG* |
| Az18 | full carboxysome, 4M CsoS2 | pHnCB | Cm | MGSNMPSQSGMNPADLSGLSGKELARARRAALSKQGKAAVSNKTASVNRSTKQAASSIN TNQVRSSVNEVPTDYQMADQLCSTIDHADFGTESNRVRDLCRQRREALSTIGKKAVKTNG KPSGRVRPQQSVVHNDAMIENAGDTNQSSSTSLNNELSEICSIADDMPERFGSQAKTVRD |

| ID | description | back-bone | anti-biotic | CsoS2 or CsoS2 fragment sequence |
|---|---|---|---|---|
| | | | | ICRARRQALSERGTRAVPPKPQSQGGPGRNGYQIDGYLDTALHGRDAAKRHREMLCQY<br>GRGTAPSCKPTGRVKNSVQSGNAAPKKVETGHTLSGGSVTGTQVDRKSHVTGNEPGTC<br>RAVTGTEYVGTEQFTSFCNTSPKPNATKVGQDRTNKGQSVTGNLVDRSELVTGNEPGSC<br>SRVTGSQYGQSKICGGGVGKVRSMRTLRGTSVSGQQLDHAPKMSGDERGGCMPVTGN<br>EYYGREHFEPFCTSTPEPEAQSTEQSLTCEGQIISGTSVDASDLVTGNEIGEQQLISGDAY<br>VGAQQTGCLPTSPRFNQTGNVQSMGFKNTNQPEQNFAPGEVMPTDFSIQTPARSAQNRI<br>TGNDIAPSGRITGPGMLATGLITGTPEFRHAARELVGSPQPMAMAMANRNKAAQAPVVQP<br>EVVATQEKPELVCAPRSDQMDRVSGEGKERCHITGDDWSVNKHITGTAGQWASGRNPS<br>MRGNARVVETSAFANRNVPKPEKPGSKITGSSGNDTQGSLITYSGGARG* |
| Az07 | full carboxysome, 13M CsoS2 | pHnCB | Cm | MGSNMPSQSGMNPADLSGLSGKELARARRAALSKQGKAAVSNKTASVNRSTKQAASSIN<br>TNQVRSSVNEVPTDYQMADQLCSTIDHADFGTESNRVRDLCRQRREALSTIGKKAVKTNG<br>KPSGRVRPQQSVVHNDAMIENAGDTNQSSSTSLNNELSEICSIADDMPERFGSQAKTVRD<br>ICRARRQALSERGTRAVPPKPQSQGGPGRNGYQIDGYLDTALHGRDAAKRHREMLCQY<br>GRGTAPSCKPTGRVKNSVQSGNAAPKKVETGHTLSGGSVTGTQVDRKSHVTGNEPGTC<br>RAVTGTEYVGTEQFTSFCNTSPKPNATKVNVTTTARGRPVSGTEVSRTEKVTGNESGVC<br>RNVTGTEYMSNEAHFSLCGTAAKPSQADKVMFGATARTHQVVSGSDEFRPSSVTGNES<br>GAKRTITGSQYADEGLARLTINGAPAKVARTHTFAGSDVTGTEIGRSTRVTGDESGSCRSI<br>SGTEYLSNEQFQSFCDTKPQRSPFKVGQDRTNKGQSVTGNLVDRSELVTGNEPGSCSRV<br>TGSQYGQSKICGGGVGKVRSMRTLRGTSVSGQQLDHAPKMSGDERGGCMPVTGNEYY<br>GREHFEPFCTSTPEPTRVKNSVQSGNAAPKKVETGHTLSGGSVTGTQVDRKSHVTGNEP<br>GTCRAVTGTEYVGTEQFTSFCNTSPKPNATKVNVTTTARGRPVSGTEVSRTEKVTGNES<br>GVCRNVTGTEYMSNEAHFSLCGTAAKPSQADKVMFGATARTHQVVSGSDEFRPSSVTG<br>NESGAKRTITGSQYADEGLARLTINGAPAKVARTHTFAGSDVTGTEIGRSTRVTGDESGSC<br>RSISGTEYLSNEQFQSFCDTKPQRSPFKVGQDRTNKGQSVTGNLVDRSELVTGNEPGSC<br>SRVTGSQYGQSKICGGGVGKVRSMRTLRGTSVSGQQLDHAPKMSGDERGGCMPVTGN<br>EYYGREHFEPFCTSTPEPEAQSTEQSLTCEGQIISGTSVDASDLVTGNEIGEQQLISGDAY<br>VGAQQTGCLPTSPRFNQTGNVQSMGFKNTNQPEQNFAPGEVMPTDFSIQTPARSAQNRI<br>TGNDIAPSGRITGPGMLATGLITGTPEFRHAARELVGSPQPMAMAMANRNKAAQAPVVQP<br>EVVATQEKPELVCAPRSDQMDRVSGEGKERCHITGDDWSVNKHITGTAGQWASGRNPS<br>MRGNARVVETSAFANRNVPKPEKPGSKITGSSGNDTQGSLITYSGGARG* |
| Lz106 | full carboxysome, double C-peptide repeats CsoS2 | pHnCB | Cm | MPSQSGMNPADLSGLSGKELARARRAALSKQGKAAVSNKTASVNRSTKQAASSINTNQV<br>RSSVNEVPTDYQMADQLCSTIDHADFGTESNRVRDLCRQRREALSTIGKKAVKTNGKPSG<br>RVRPQQSVVHNDAMIENAGDTNQSSSTSLNNELSEICSIADDMPERFGSQAKTVRDICRA<br>RRQALSERGTRAVPPKPQSQGGPGRNGYQIDGYLDTALHGRDAAKRHREMLCQYGRGT |

| ID | description | back-bone | anti-biotic | CsoS2 or CsoS2 fragment sequence |
|---|---|---|---|---|
| | | | | APSCKPTGRVKNSVQSGNAAPKKVETGHTLSGGSVTGTQVDRKSHVTGNEPGTCRAVTGTEYVGTEQFTSFCNTSPKPNATKVNVTTTARGRPVSGTEVSRTEKVTGNESGVCRNVTGTEYMSNEAHFSLCGTAAKPSQADKVMFGATARTHQVVSGSDEFRPSSVTGNESGAKRTITGSQYADEGLARLTINGAPAKVARTHTFAGSDVTGTEIGRSTRVTGDESGSCRSISGTEYLSNEQFQSFCDTKPQRSPFKVGQDRTNKGQSVTGNLVDRSELVTGNEPGSCSRVTGSQYGQSKICGGGVGKVRSMRTLRGTSVSGQQLDHAPKMSGDERGGCMPVTGNEYYGREHFEPFCTSTPEPEAQSTEQSLTCEGQIISGTSVDASDLVTGNEIGEQQLISGDAYVGAQQTGCLPTSPRFNQTGNVQSMGFKNTNQPEQNFAPGEVMPTDFSIQTPARSAQNRITGNDIAPSGRITGPGMLATGLITGTPEFRHAARELVGSPQPMAMAMANRNKAAQAPVVQPEVVATQEKPELVCAPRSDQMDRVSGEGKERCHITGDDWSVNKHITGTAGQWASGRNPSMRGNATNQPEQNFAPGEVMPTDFSIQTPARSAQNRITGNDIAPSGRITGPGMLATGLITGTPEFRHAARELVGSPQPMAMAMANRNKAAQAPVVQPEVVATQEKPELVCAPRSDQMDRVSGEGKERCHITGDDWSVNKHITGTAGQWASGRNPSMRGNARVVETSAFANRNVPKPEKPGSKITGSSGNDTQGSLITYSGGARG* |
| Lz105 | full carboxysome, double CTD CsoS2 | pHnCB | Cm | MPSQSGMNPADLSGLSGKELARARRAALSKQGKAAVSNKTASVNRSTKQAASSINTNQVRSSVNEVPTDYQMADQLCSTIDHADFGTESNRVRDLCRQRREALSTIGKKAVKTNGKPSGRVRPQQSVVHNDAMIENAGDTNQSSSTSLNNELSEICSIADDMPERFGSQAKTVRDICRARRQALSERGTRAVPPKPQSQGGPGRNGYQIDGYLDTALHGRDAAKRHREMLCQYGRGTAPSCKPTGRVKNSVQSGNAAPKKVETGHTLSGGSVTGTQVDRKSHVTGNEPGTCRAVTGTEYVGTEQFTSFCNTSPKPNATKVNVTTTARGRPVSGTEVSRTEKVTGNESGVCRNVTGTEYMSNEAHFSLCGTAAKPSQADKVMFGATARTHQVVSGSDEFRPSSVTGNESGAKRTITGSQYADEGLARLTINGAPAKVARTHTFAGSDVTGTEIGRSTRVTGDESGSCRSISGTEYLSNEQFQSFCDTKPQRSPFKVGQDRTNKGQSVTGNLVDRSELVTGNEPGSCSRVTGSQYGQSKICGGGVGKVRSMRTLRGTSVSGQQLDHAPKMSGDERGGCMPVTGNEYYGREHFEPFCTSTPEPEAQSTEQSLTCEGQIISGTSVDASDLVTGNEIGEQQLISGDAYVGAQQTGCLPTSPRFNQTGNVQSMGFKNTNQPEQNFAPGEVMPTDFSIQTPARSAQNRITGNDIAPSGRITGPGMLATGLITGTPEFRHAARELVGSPQPMAMAMANRNKAAQAPVVQPEVVATQEKPELVCAPRSDQMDRVSGEGKERCHITGDDWSVNKHITGTAGQWASGRNPSMRGNARVVETSAFANRNVPKPEKPGSKITGSSGNDTQGSLITYSGGARGTNQPEQNFAPGEVMPTDFSIQTPARSAQNRITGNDIAPSGRITGPGMLATGLITGTPEFRHAARELVGSPQPMAMAMANRNKAAQAPVVQPEVVATQEKPELVCAPRSDQMDRVSGEGKERCHITGDDWSVNKHITGTAGQWASGRNPSMRGNARVVETSAFANRNVPKPEKPGSKITGSSGNDTQGSLITYSGGARG* |
| Az25 | 1M-GFP | pFA | Kan | MSGRVKNSVQSGNAAPKKVETGHTLSGGSVTGTQVDRKSHVTGNEPGTCRAVTGTEYV |

| ID | description | back-bone | anti-biotic | CsoS2 or CsoS2 fragment sequence |
| --- | --- | --- | --- | --- |
| | | | | GTEQFTSFCNTSPTVSKGEELFTGVVPILVELDGDVNGHKFSVRGEGEGDATNGKLTLKFI<br>CTTGKLPVPWPTLVTTLTYGVQCFSRYPDHMKQHDFFKSAMPEGYVQERTISFKDDGTY<br>KTRAEVKFEGDTLVNRIELKGIDFKEDGNILGHKLEYNFNSHNVYITADKQKNGIKANFKIRH<br>NVEDGSVQLADHYQQNTPIGDGPVLLPDNHYLSTQSKLSKDPNEKRDHMVLLEFVTAAGI<br>TLGMDELYKSGHHHHHH* |
| Az26 | 2M-GFP | pFA | Kan | MSGRVKNSVQSGNAAPKKVETGHTLSGGSVTGTQVDRKSHVTGNEPGTCRAVTGTEYV<br>GTEQFTSFCNTSPKPNATKVNVTTTARGRPVSGTEVSRTEKVTGNESGVCRNVTGTEYM<br>SNEAHFSLCGTAATVSKGEELFTGVVPILVELDGDVNGHKFSVRGEGEGDATNGKLTLKFI<br>CTTGKLPVPWPTLVTTLTYGVQCFSRYPDHMKQHDFFKSAMPEGYVQERTISFKDDGTY<br>KTRAEVKFEGDTLVNRIELKGIDFKEDGNILGHKLEYNFNSHNVYITADKQKNGIKANFKIRH<br>NVEDGSVQLADHYQQNTPIGDGPVLLPDNHYLSTQSKLSKDPNEKRDHMVLLEFVTAAGI<br>TLGMDELYKSGHHHHHH* |
| Az27 | 4M-GFP | pFA | Kan | MSGRVKNSVQSGNAAPKKVETGHTLSGGSVTGTQVDRKSHVTGNEPGTCRAVTGTEYV<br>GTEQFTSFCNTSPKPNATKVNVTTTARGRPVSGTEVSRTEKVTGNESGVCRNVTGTEYM<br>SNEAHFSLCGTAAKPSQADKVMFGATARTHQVVSGSDEFRPSSVTGNESGAKRTITGSQ<br>YADEGLARLTINGAPAKVARTHTFAGSDVTGTEIGRSTRVTGDESGSCRSISGTEYLSNEQ<br>FQSFCDTVSKGEELFTGVVPILVELDGDVNGHKFSVRGEGEGDATNGKLTLKFICTTGKLP<br>VPWPTLVTTLTYGVQCFSRYPDHMKQHDFFKSAMPEGYVQERTISFKDDGTYKTRAEVK<br>FEGDTLVNRIELKGIDFKEDGNILGHKLEYNFNSHNVYITADKQKNGIKANFKIRHNVEDGS<br>VQLADHYQQNTPIGDGPVLLPDNHYLSTQSKLSKDPNEKRDHMVLLEFVTAAGITLGMDE<br>LYKSGHHHHHH* |
| Az24 | MR (6M)-GFP | pFA | Kan | MSGRVKNSVQSGNAAPKKVETGHTLSGGSVTGTQVDRKSHVTGNEPGTCRAVTGTEYV<br>GTEQFTSFCNTSPKPNATKVNVTTTARGRPVSGTEVSRTEKVTGNESGVCRNVTGTEYM<br>SNEAHFSLCGTAAKPSQADKVMFGATARTHQVVSGSDEFRPSSVTGNESGAKRTITGSQ<br>YADEGLARLTINGAPAKVARTHTFAGSDVTGTEIGRSTRVTGDESGSCRSISGTEYLSNEQ<br>FQSFCDTKPQRSPFKVGQDRTNKGQSVTGNLVDRSELVTGNEPGSCSRVTGSQYGQSKI<br>CGGGVGKVRSMRTLRGTSVSGQQLDHAPKMSGDERGGCMPVTGNEYYGREHFEPFCT<br>STPEPTVSKGEELFTGVVPILVELDGDVNGHKFSVRGEGEGDATNGKLTLKFICTTGKLPV<br>PWPTLVTTLTYGVQCFSRYPDHMKQHDFFKSAMPEGYVQERTISFKDDGTYKTRAEVKF<br>EGDTLVNRIELKGIDFKEDGNILGHKLEYNFNSHNVYITADKQKNGIKANFKIRHNVEDGSV<br>QLADHYQQNTPIGDGPVLLPDNHYLSTQSKLSKDPNEKRDHMVLLEFVTAAGITLGMDEL<br>YKSGHHHHHH* |
